# A rapid method to quantify vein density in C_4_ plants using starch staining

**DOI:** 10.1101/2023.03.06.531345

**Authors:** Conor J. C. Simpson, Pallavi Singh, Deedi E.O. Sogbohossou, M. Eric Schranz, Julian M. Hibberd

## Abstract

C_4_ photosynthesis has evolved multiple times in the angiosperms and typically involves alterations to the biochemistry, cell biology and development of leaves. One common modification found in C_4_ plants compared with the ancestral C_3_ state is an increase in vein density such that the leaf contains a larger proportion of bundle sheath cells. Recent findings indicate that there may be significant intra-specific variation in traits such as vein density in C_4_ plants but to use such natural variation for trait-mapping, rapid phenotyping would be required. Here we report a high-throughput method to quantify vein density that leverages the bundle sheath specific accumulation of starch found in C_4_ species. Starch staining allowed high-contrast images to be acquired that permitted image analysis using a MATLAB-based program. The method works for the dicotyledon *Gynandropsis gynandra* where significant variation in vein density was detected between natural accessions, and the monocotyledon *Zea mays* where no variation was apparent in the genotypically diverse lines assessed. We anticipate this approach will be useful to map genes controlling vein density in C_4_ species demonstrating natural variation for this trait.

**One sentence summary:** Preferential accumulation of starch in bundle sheath cells of C_4_ plants allows high-throughput phenotyping of vein density.

## Introduction

Photosynthesis is the basis of life on Earth and central to this process is the enzyme Ribulose 1,5-Bishosphate Carboxylase/Oxygenase (RuBisCO) that operates in the Calvin-Benson-Bassham cycle to fix CO_2_. However, oxygenase activity RuBisCO results in the energy-intensive photorespiratory pathway (Driever and Kromdijk, 2013) and in some environments costs of photorespiration are thought to have driven the evolution of carbon concentrating mechanisms such as C_4_ photosynthesis and Crassulacean Acid Metabolism (CAM). Thus, C_3_ photosynthesis is considered the ancestral state and despite its complexity, derived states involving carbon concentrating mechanisms are thought to have evolved repeatedly (Edwards, 2019). In the case of C_4_ photosynthesis, current estimates are that it has evolved independently in more than sixty-six lineages (Sage *et al*., 2011). The vast majority of C_4_ plants evolved specialized anatomy to compartmentalise photosynthesis between two cell types. Typically, this involves the mesophyll being the site of initial HCO_3_^-^ assimilation by Phospho*enol*pyruvate Carboxylase (PEPC) and the bundle sheath being repurposed for the Calvin-Benson-Bassham cycle. Compared with the C_3_ state, the partitioning of photosynthesis between these two cell types is associated with changes to transcriptional, post-transcriptional and also post-translational regulatory mechanisms (Hibberd and Covshoff, 2010; Reeves *et al*., 2017). For some genes encoding components of the C_4_ cycle a detailed understanding has emerged such that changes in *cis* (Nomura *et al*., 2000, *2005a,b;* Gowik *et al*., 2004, 2017; Williams *et al*., 2016) or *trans* (Brown *et al*., 2011; Reyna-Llorens *et al*., 2018) are considered the driving forces behind alterations to gene expression. In contrast, we have a more limited understanding of the genes underpinning the evolution of Kranz anatomy and how their expression has changed during the C_3_ to C_4_ transition (Sedelnikova *et al*., 2018).

Reports of significant natural variation in traits including components of Kranz anatomy (Lundgren *et al*., 2016; Yabiku and Ueno, 2017; Kolbe and Cousins, 2018; Reeves *et al*., 2018) provide an opportunity to probe the basis of this trait using quantitative genetics. For example, if it were possible to screen vein density or bundle sheath size rapidly in populations made up of genetically diverse individuals such as Multi-Advanced Generation Inter-Crossing (MAGIC) populations or natural accessions designed for Genome Wide Association Study (GWAS) it should be possible to start to link genes with Kranz anatomy. This approach would be facilitated by rapid phenotyping of these traits. Significant advances have been made in phenotyping at scale for protein activity assays (Gibon *et al*., 2004; Sulpice *et al*., 2010) or the use of robotics for data collection in the field (Virlet *et al*., 2017). Notably, using a phenotyping facility Guo *et al*. (2018) rapidly analyzed fifty-one image-based traits in rice under drought stress. Correlating these traits with traditional drought resistance traits enabled novel genes linked to drought resistance to be identified via a GWAS. Such approaches may need support from non-invasive imaging techniques (Yang *et al*., 2020) and consequently high-throughput image processing is key to alleviate the phenotyping bottleneck associated with laborious, error-prone manual image analysis (Moen *et al*., 2019). Image analysis using Convoluted Neural Networks (CNNs) has been used to map Quantitative Trait Loci (QTL) associated with stomata size and density in maize (Xie *et al*., 2021). Moreover, image segmentation tools have been developed to quantify vein density (Dhondt *et al*., 2012; Parsons-Wingerter *et al*., 2014; Bühler *et al*., 2015), but such automated approaches have not yet replaced manual tracing (Perez-Harguindeguy *et al*., 2016). Moreover, analysis of tissues such as cotyledons of *Arabidopsis thaliana* (Dhondt *et al*., 2012; Parsons-Wingerter *et al*., 2014) can require extensive manual supervision and may not be suited for tissues with higher vein density. Although, the powerful programme LeafVeinCNN (Xu *et al*., 2021) enables high-throughput analysis of traits including vein order, areole number, vein width and vein density it currently requires manual real-time supervision of individual images and can be computationally slow due to the number of networks in use and the number of measurements being attained per image (Xu *et al*., 2021). Further, to our knowledge, only one tool has been developed to assess vein structure in monocotyledons (Robil *et al*., 2021). As C_4_ photosynthesis is found in both monocotyledons and dicotyledons it would be useful to have a tool that permits analysis of vein density in both clades.

Here we aimed to test whether preferential accumulation of starch in the bundle sheath of C_4_ plants (Lunn and Furbank, 1997; Figure 1A) could be used to generate high-contrast images so that a segmentation-based method could rapidly assess vein density in these plants. We show that such an approach can be used to measure vein density in both monocotyledons and dicotyledons. Moreover, this allowed us to develop a computational pipeline enabling high-contrast black and white images to be processed automatically using the image processing toolbox in MATLAB. Not only does the method allow rapid phenotyping but it also showed that whilst in genotypically diverse founders of a maize population (Dell’Acqua *et al*., 2015) there was little variation in vein density, in the C_4_ dicotyledenous model *Gynandropsis gynandra* there is significant natural variation in vein density.

**Figure 1:**
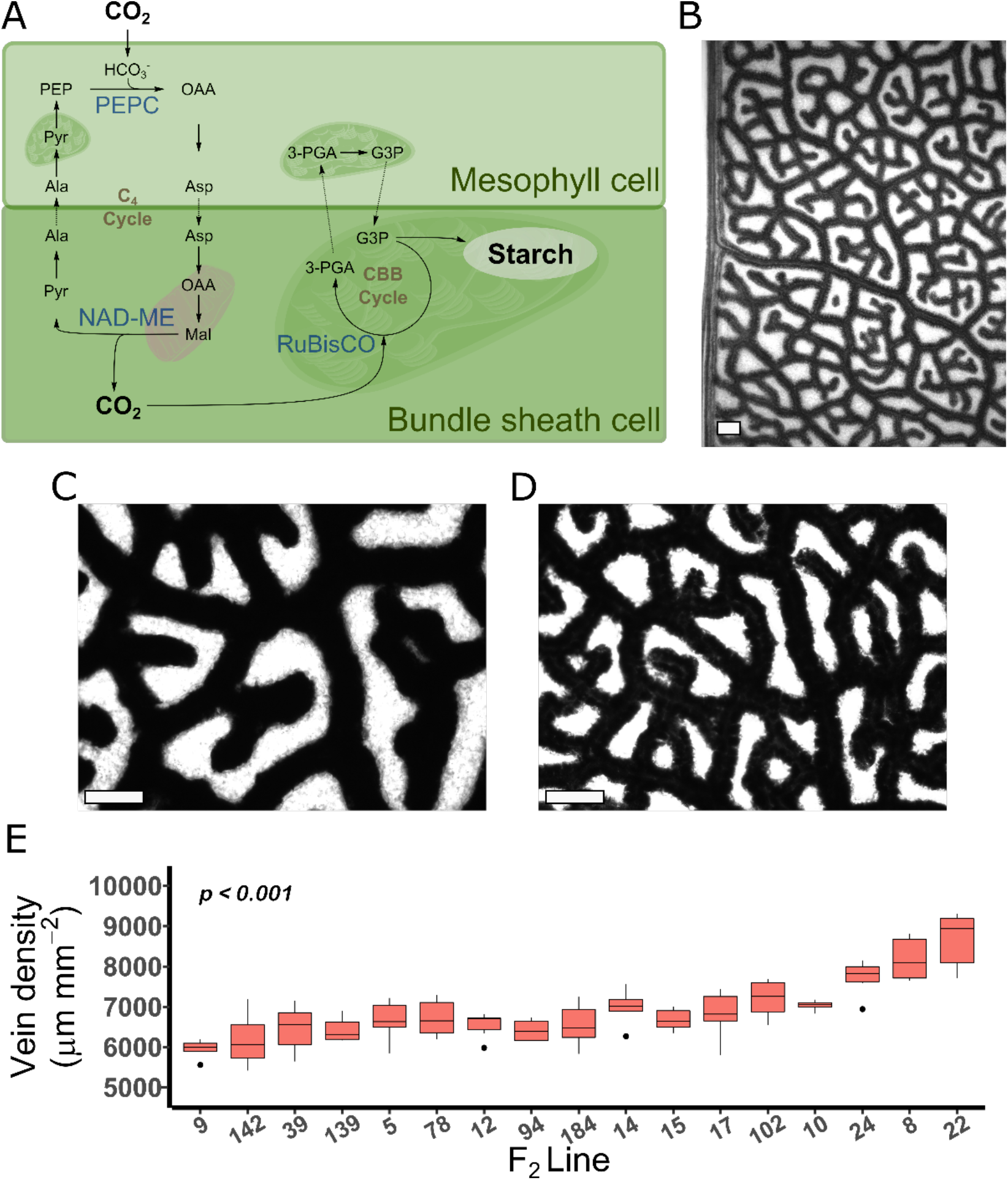
Starch staining in the bundle sheath of C_4_ *Gynandropsis gynandra* demonstrates natural variation in vein density across a diversity panel. **(A)** Schematic of the C_4_ pathway. Arrows indicate carbon flux. Enzymes involved in carboxylation and decarboxylation shown in blue. Intracellular compartments represented schematically with mitochondria in pink and chloroplasts dark green. **(B, C, D)** Representative images of *Gynandropsis gynandra* leaves stained for starch at low magnification **(B)**, higher magnification with low vein density **(C)** and higher magnification with high vein density **(D). (E)** Natural variation in vein density across seventeen lines. Abbreviations as follows: PEP = Phospho*enol*pyruvate; OAA = Oxaloacetate; Asp = Aspartate; Mal = Malate; Pyr = Pyruvate; Ala = Alanine; 3-PGA = 3-Phosphoglycerate; G3P = Glyceraldehyde 3-phosphate; PEPC = Phospho*enol*pyruvate carboxylase; NAD-ME = NAD-Malic Enzyme; RuBisCO = Ribulose Bisphosphate Carboxylase/Oxygenase; CBB = Calvin Benson Bassham cycle. Scale bars represent 200μm. Six images per plant were assessed and p-value calculated by ANOVA.

## Results and Discussion

### Natural variation for vein density in C_4_ *G. gynandra* and an automated image processing tool

Staining leaves of *G. gynandra* clearly indicated preferential accumulation of starch in the bundle sheath (Figure 1B). We therefore subjected an F_2_ population of *G. gynandra* comprising 199 lines derived from a cross between parents differing in vein density (Reeves *et al*., 2018) to starch staining. Initially, a random subset of seventeen plants were stained and this indicated significant natural variation in vein density amongst the F_2_ plants (*p* < 0.001; Figure 1C,D&E). We reasoned that this variance in vein density between accessions provided a resource to develop and ground-truth a semi-automated procedure to quantify vein density.

Using MATLAB’s Image Processing Toolbox (2019a) an automated pipeline, hereafter Starch4Kranz, was developed to reliably quantify bundle sheath density. This pipeline involved converting each image to greyscale (Figure 2A), enhancing the contrast to improve differences between bundle sheath strands and the background (Figure 2B) and then blurring such that a given pixel is converted to the mean pixel intensity in a sub-matrix of 2w+1 where w is 20. Thus, for each pixel its value is converted to the average of the neighboring sub-matrix of 20×20 pixels. This smoothing resulted in pixels in each bundle sheath strand having similar values, and the same was true for background pixels (Figure 2C). Images are then binarised (Figure 2D) such that stained regions appear white (with a pixel value of 1) and the background is black (with a pixel value of 0). Skeletonisation converts all pixels with a value of 1 to curved lines a single pixel in width (Figure 2E). Lastly, these skeletonised images are trimmed such that superfluous pixels are removed (Figure 2F, Figure S1). The number of pixels trimmed is defined by the user as the trim_factor, but if the program detects a significantly larger number of superfluous pixels compared with expected vein density then the trim_factor can be calculated automatically based on the mode length of connected superfluous pixels. At this stage bundle sheath strands are represented by a line that is one pixel thick and so the total number of pixels provides an estimate of vein length. Vein density can then be determined from the total number of pixels and the image size.

**Figure 2:**
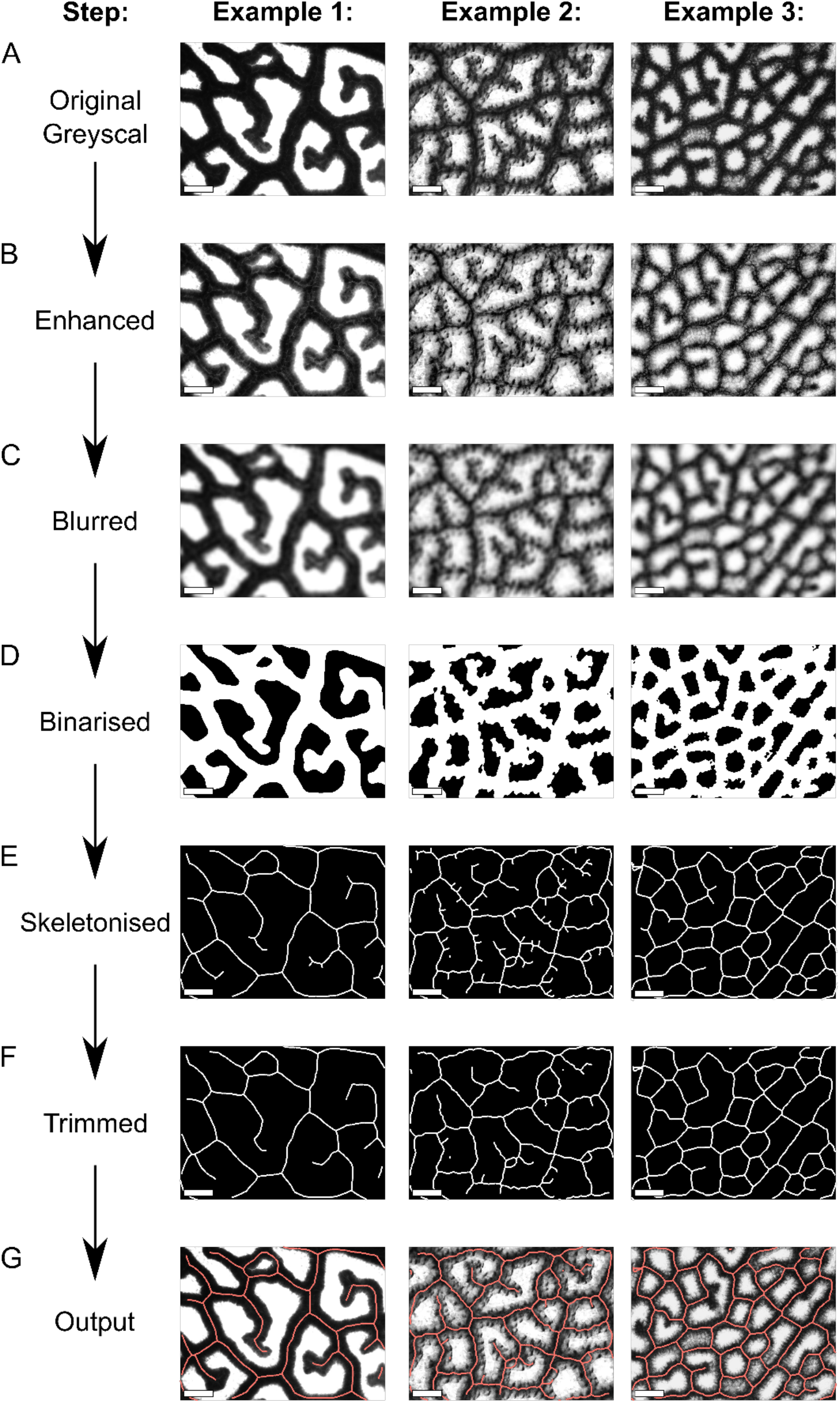
Summary of critical automation steps. **(A)** A greyscale image is used as input and subjected to contrast enhancement **(B),** followed by blurring **(C)**, binarization **(D)**, skeletonization **(E)**, and trimming **(F)**. An output is then produced for the user to check **(G)**. Scale bars represent 200μm. Skeletons have been thickened to ensure visibility.

Starch4Kranz processes all images in a given directory and saves results into a table containing image name and vein density. After each image is processed a window appears containing the vein trace superimposed on the original image in greyscale (Figure 2G). Compared with approximately eleven hours reuired to manually trace around 102 images (Figure 1E) implementing the Starch4Kranz pipeline for the same samples took fifteen minutes.

### Automatic quantification of high contrast images for vein density

The Starch4Kranz pipeline was next implemented on data from the seventeen F_2_ plants for which vein density had been determined manually. From 102 images, the same pattern of variation in vein density was observed (Figure 3A) and estimates from manual and automated pipelines were highly correlated (*r* = 0.92; Figure 3B). When Starch4Kranz was applied to 1101 images, 98.5% passed the manual check. Of the seventeen images that did not, manual inspection indicated they were either poorly stained or images were of poor quality. When twenty-seven images were subjected to the LeafVeinCNN pipeline (Xu *et al*., 2021) three and a half minutes were required to run the CNNs but repeated manual input was required to obtain optimal thresholding. Moreover, compared with LeafVeinCNN (*r* = 0.74; Figure S2B) the outputs from Starch4Kranz pipeline correlated more strongly with manual tracings (*r* = 0.95; Figure S2A). LeafVeinCNN performed less well when vein densities were low (Figure S2C), which may be because it was trained on data derived from leaves imaged at lower magnification.

**Figure 3:**
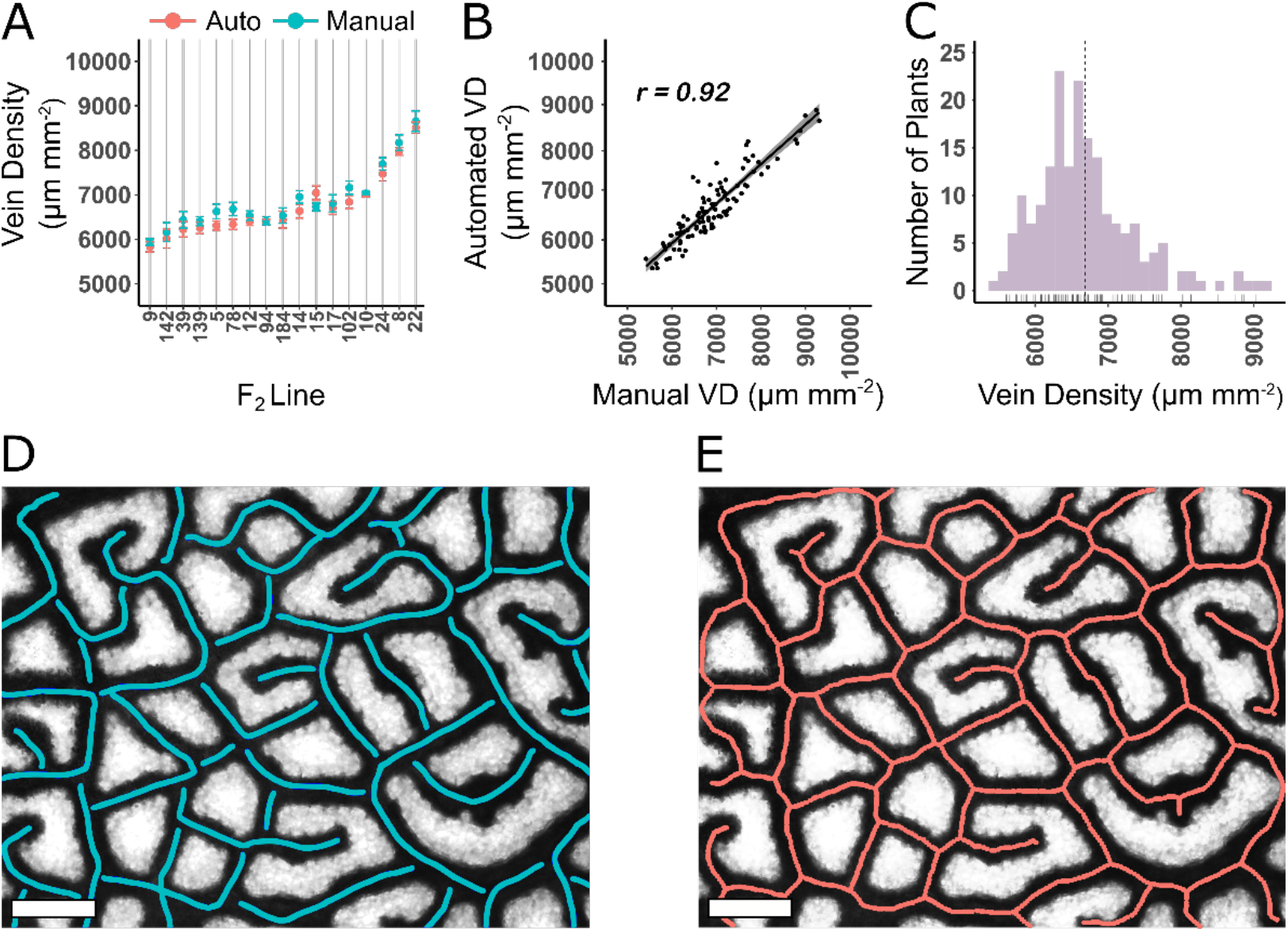
Ground truthing of automated analysis. **(A)** Seventeen randomly selected lines from the 199 line population were assessed both automatically (red) and manually (blue) for vein density. **(B)** Correlation analysis between automatic and manual quantification of vein density. VD = Vein Density. Line of best fit shown with grey shading representing 95% confidence intervals. *r* = Pearson’s correlation coefficient. **(C)** Distribution of vein density across 199 plants determined using the automated pipeline Starch4Kranz. **(D)** Example of a manually traced image. **(E)** Example of the automated output. Scale bars represent 200*μm*. Skeletons have been thickened to ensure visibility.

We found that subdividing an image into multiple sections, and capturing slightly out-of-focus images could enhance contrast between veins and background. This was of particular use for samples with high vein densities, or when staining was less good. As we found that some images did not allow accurate vein density to be determined, a manual check is incorporated into the pipeline (Figure 3B). Compared with other methods that require continual manual checking of images during processing (Dhondt *et al*., 2012; Parsons-Wingerter *et al*., 2014; Bühler *et al*., 2015; Xu *et al*., 2021) Starch4Kranz saves time because it runs in the background across multiple images, and a manual check does not take place until the programme has finished. Machine learning techniques for image analysis are a useful tool for quantitative studies in plant sciences and have been employed to identify QTL associated with stomatal density in maize (Xie *et al*., 2021) and estimate heritability of vein density in sunflower (Earley *et al*., 2022). However, such approaches can require extensive computational understanding as well as power to build neural networks, and depending on the complexity of the phenotype may require manual annotation of thousands of images for network training (Moen *et al*., 2019; van Dijk *et al*., 2021). Thus, LeafVeinCNN is a powerful tool suitable for quantifying multiple traits associated with vein architecture, but Starch4Kranz offers a rapid and robust tool allowing vein density to be detrmined in C_4_ species. It also confirmed significant natural variation amongst F_2_ lines of *G. gynandra* (Figure 3C).

### Natural variation in vein density was not detectable in founder lines of a maize MAGIC population

Aside from Grasviq (Robil *et al*., 2021) we noted a lack of automated methods for the assessment of vein density in monocotyledons. Because the preferential accumulation of starch in bundle sheath cells had first been reported in monocotyledons (Lunn and Furbank, 1997) we next assessed whether the Starch4Kranz algorithm could be used in maize. For this, an additional step was added to the pipeline to remove commissural veins (Figure S3), which are not considered important components of Kranz anatomy (Langdale, 2011). We also noted that they stained for starch inconsistently likely because they are considered a simplified vascular tissue not encircled by bundle sheath cells (Sakaguchi and Fukuda, 2008; Sakaguchi *et al*., 2010). If commissural veins are of interest, and sufficient contrast obtained, then “auto” could be selected as an input variable instead of “monocot” (see Methods) and commissural veins will not be removed. Commissural veins were defined as the shortest branch stemming from a pixel with three branch points. Pattern recognition allowed these to be identified and removed (Figure S3). To ground-truth this approach, manual tracings that did not include these veins were undertaken. Images from starch staining of nine founder lines of a MAGIC population from maize (Dell’Acqua *et al*., 2015) were subjected to Starch4Kranz. Estimates were highly correlated with manual analysis (r = 0.91; Figure 4A) but no statistically significant differences were observed between genotypes (*p* > 0.05; Figure 4B). Thus, Starch4Kranz can be used to quantify vein density in C_4_ monocotyledons in addition to dicotyledons (Figure 4&S4). However, for the founder lines studied here, we detected no significant differences in vein density (Figure 4B).

**Figure 4:**
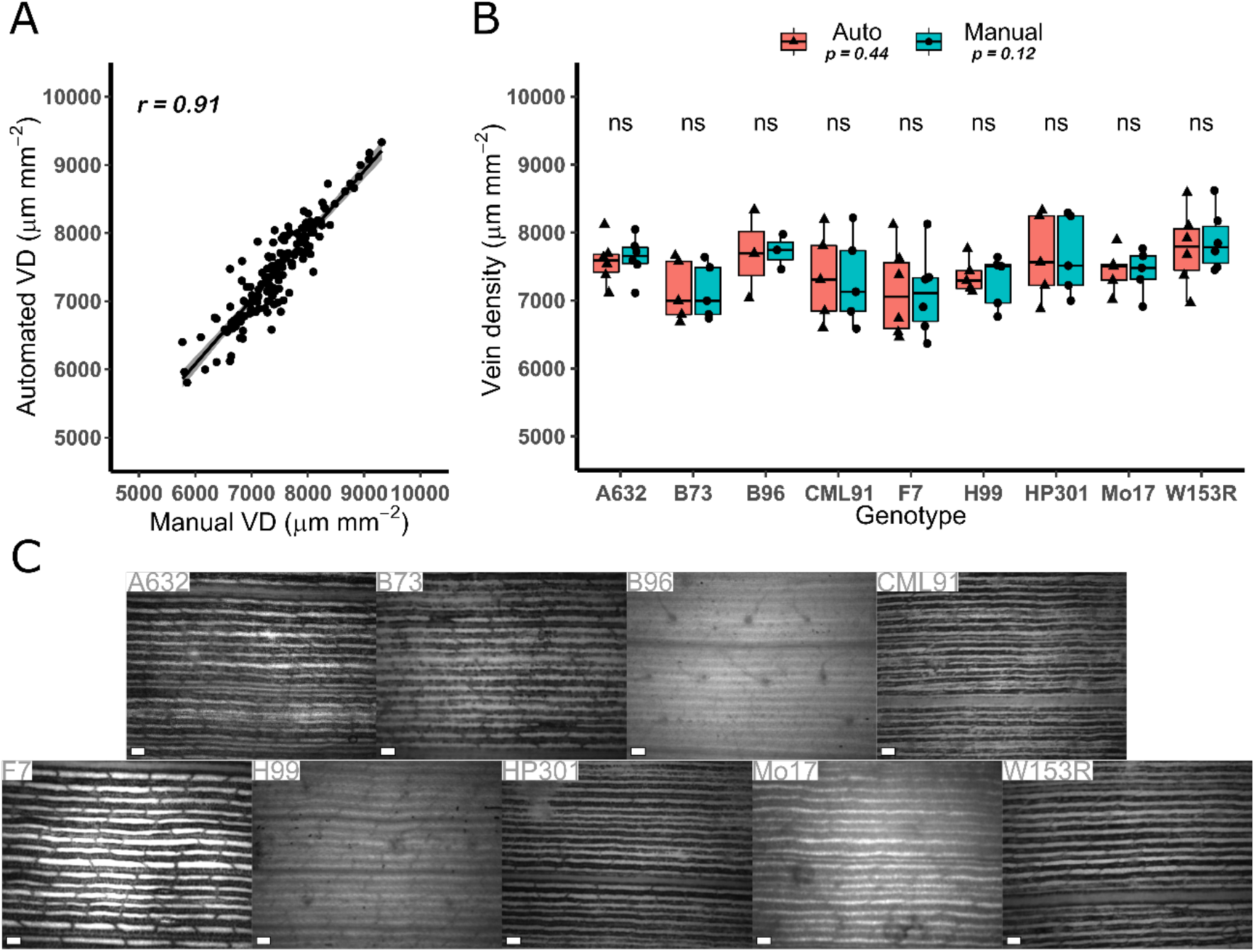
Starch staining of vein density in founder lines of a maize MAGIC population. **(A)** Vein density in maize determined automatically and manually after starch staining. **(B)** No significant differences between founder lines of maize were detected from either manual (red) or automatic (blue) methods. r = Pearson’s correlation coefficient; VD = Vein Density; p-values determined by ANOVA and student’s t-test. **(C)** Representative images of maize leaves stained for starch. Scale bars represent 200μm.

This finding emphasises the need to generate specific mapping populations designed for the trait of interest. *G. gynandra* F_2_ lines produced from a cross between two lines phenotypically distinct for vein density displayed abundant natural variation in this trait (Figures 3C). Although variation in vein density has been reported in maize (Kolbe and Cousins, 2018), equivalent variation was not present in the MAGIC maize founders (Figure 4B) indicating that the MAGIC maize population (Dell’Acqua *et al*., 2015) is unlikely to be suitable for studying QTL underlying vein density. That said, the ease and speed with which Starch4Kranz permits quantification of vein density means that screening other C_4_ mapping populations such NAM populations of maize (Yu *et al*., 2008; McMullen *et al*., 2009) or MAGIC populations of sorghum (Ongom and Ejeta, 2017) could be useful.

In summary, we report a simple protocol allowing vein density to be determined from starch staining of vascular tissue in C_4_ species. Automatation of this pipeline allowed rapid and accurate quantification of vein density. As increased vein density is considered an important trait allowing C_4_ photosynthesis (Sage, 2001), where intra-specific natural variation for this trait exists our approach could provide insight into mechanisms evolution has co-opted to generate the complex C_4_ phenotype.

## Methods

### Plant Growth and leaf sampling

*Gynandropsis gynandra* was grown under irrigated glasshouse conditions from April to July 2019. Temperatures were maintained at 24°C in the day and 20°C during the night, with artificial lighting maintaining a minimum light intensity of 300 *μmol m^-2^s^-1^* and a photoperiod of 16 hours light and 8 hours dark. Relative humidity was 60%. Maize was grown under field conditions from April to August 2022 at NIAB, Cambridge UK. Plants were treated with pre-emptive herbicide (Stomp^®^ Aqua) one day after sowing, fertilizer (ammonium nitrate) five days after sowing, molluscicide (Sluxx) one month after sowing, and herbicide (Leystar^®^) two months after sowing. During the growing season, average temperature was 19.8°C during the day and 15.2°C during the night. Maximum and minimum daytime temperature was 39.9°C and 8.9°C and that during the night was 34.2°C and 2.9°C. Total precipitation was 89mm and average relative humidity was 65%.

*G. gynandra* leaves were harvested over a three-day period from plants four weeks after germination. The youngest fully expanded leaf was selected. Tissue was harvested from the central leaflet of each leaf and immediately submerged in a plastic cuvette containing 3:1 100% (v/v) ethanol: acetic acid solution for 4 hours before treatment with 70% (v/v) ethanol overnight at 37°C, being refreshed one hour into treatment. Samples were then cleared using 5% (w/v) NaOH for 3 hours at 37°C before being washed with, and replaced in, 70% (v/v) ethanol until preparation for imaging. Maize leaves were harvested over a two-week period as part of a larger field trial. All disks were taken from the most recently fully expanded leaf. Leaves were harvested in the field and kept in water until leaf disks were taken. One disk was taken per leaf and immediately submerged in 3:1 100% (v/v) ethanol:acetic acid fixative solution for 4 hours before being transferred to 70% (v/v) ethanol solution and left overnight at 37°C. Samples were then cleared using 5% (w/v) NaOH for 6 hours at 37°C before being washed with, and replaced in 70% (v/v) ethanol until preparation for imaging.

Prior to imaging, leaf disks were placed in Lugol’s solution. Specifically, using forceps each disk was transferred to Lugol’s solution and then removed immediately after submersion and placed int 70% (v/v) ethanol to minimize stain accumulation. If the stain did not accumulate rapidly the sample was re-placed in Lugol’s solution until it stain had been taken up. Excess Lugol’s solution was washed off in 70% (v/v) ethanol. Each leaf disk was then mounted on a slide using water and imaged at a magnification of 100x on an Olympus BX41 light microscope with a mounted Micropublisher 3.3 RTV camera (Q Imaging). Images were captured with Q-Capture Pro 7 software.

### Manual and automated analysis of vein density and statistical analysis

Veins were measured directly in ImageJ (Rasband, 2014) using the paint brush tool to quantify total length of veins in *μm*. Lengths were converted to density (*μm mm*^-2^) by dividing by the image’s area in *mm^2^*. Table 1 sumarises the important MATLAB functions and their function. In addition to this, pattern recognition was used to identify commissural veins in maize (Figure S3). From a subset of 102 images, twenty-seven were also analysed using LeafVeinCNN (Xu *et al*., 2021). These twenty-seven images comprised three images per plant from nine plants. LeafVeinCNN was run using default parameters consisting of 3 ensemble CNNs. Appropriate threshold values were determined for each image to attain optimal vein traces through segmentation. To achieve results comparable with Starch4Kranz the output value “VTotL” meaning total vein length in *mm*, was converted to *μm mm*^-2^.

**Table 1.**
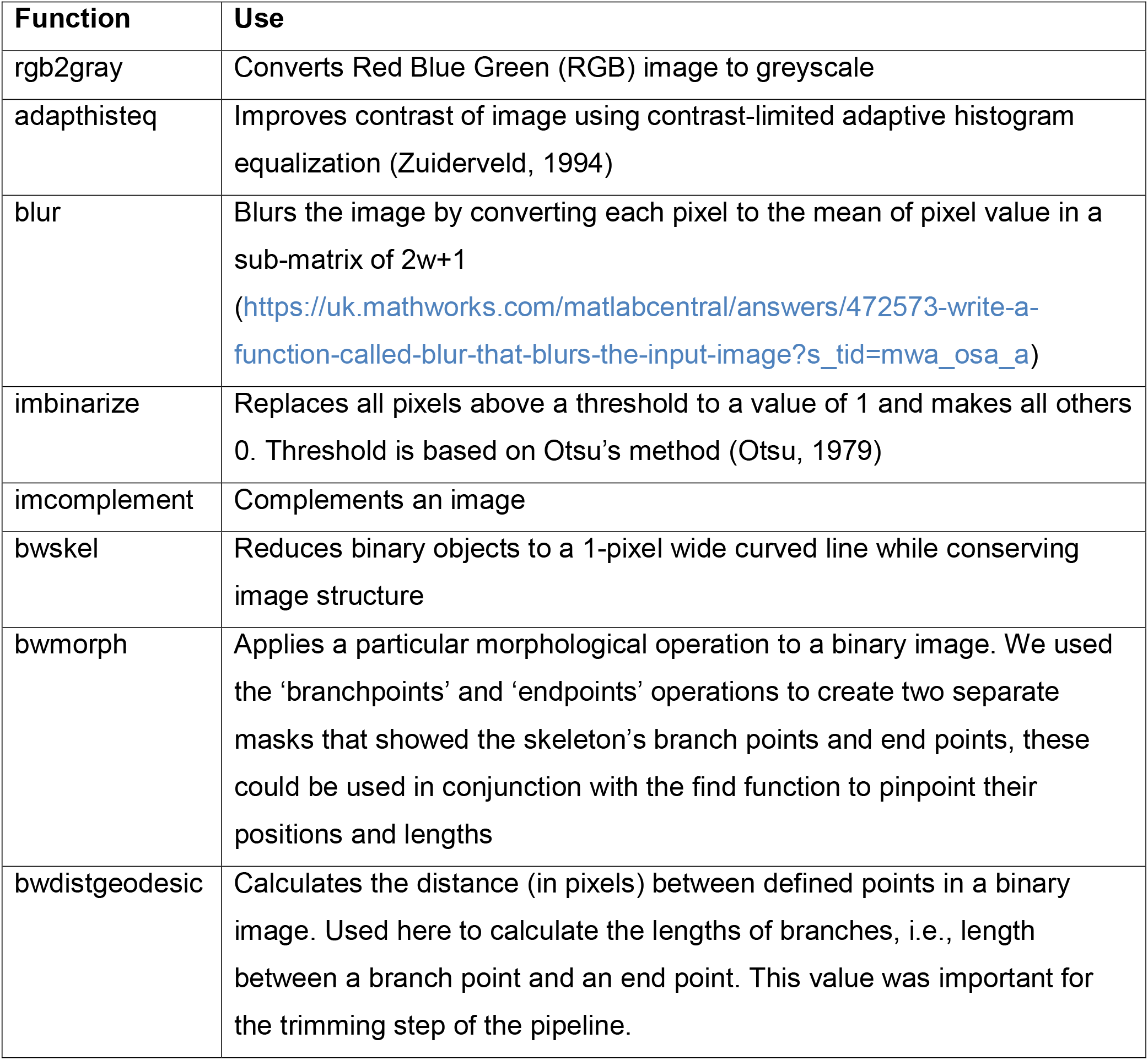
Main MATLAB base and Image Processing Toolbox functions used in Starch4Kranz.

All statistical analysis were carried out in R Studio (V: 4.1.3). Figures were generated using the ggpubr (Kassambara, 2020) package. To compare manual and automated quantification methods in *G. gynandra* a subset of seenteen plants from a total of 199 were selected randomly. Six images were assessed per plant such that the dataset was made up of 102 images. To attain a vein density measurement for each of the seventeen lines, the mean of these six images was determined. We noted that using three images yielded no significant differences in plant vein densities when comparing manual (t-test, *p* = 0.90) and automated (t-test, *p* = 0.85) methods. Therefore for maize, six replicates were used for A632, F7, and W153R; five for B73, CML91, H99, HP301, and Mo17; and three for B96. The total dataset of 138 images was assessed both manually and automatically. For both treatments all lines were normally distributed (Shapiro-Wilk’s test, *p* > 0.05) and Bartlett’s test revealed homogeneity of variance (*p* > 0.05) meaning parametric analysis was carried by Analysis Of Variance (ANOVA). Correlation analysis was carried out using the Pearson correlation coefficient. In total a *G. gynandra* population of 199 plants, each with five or six image replicates meaning a total of 1102 images were assessed for variation.

## Raw script availability and code implementation

The Starch4Kranz pipeline was designed to be simple to use and can run in the background while other tasks take place. While it requires a manual check, this can be done rapidly (~5-10 seconds per image) as the user checks each image within a whole directory once the script has stopped running. Only three scripts are needed to run the full pipeline, Starch4Kranz.m contains all executable code used in the production of our pipeline, the function, blur.m contains the blurring function, and running_Starch4Kranz.m contains a short script that runs Starch4Kranz.m while saving results to a table easily transferable to software for further data analysis. Script is available at https://github.com/plycs5/Starch4Kranz. To run the script the user must have the ImageProcessingToolbox activated and supply four inputs, filename, trim_factor (see Results), pixel_length_um (the image scale), y_pixels and x_pixels (image dimensions). A fifth input named split_mode must either be “auto”, “whole”, “split”, or “monocot”.

## ACKNOWLEDGMENTS

We thank Angie Burnett, John Ferguson, and Johannes Kromdijk for provision of maize leaf material. The work was supported by a BBSRC DTP studentship to CJCS, European Research Council Grant 694733 Revolution to JMH, and Netherlands Organization for Scientific Research Grant W.08.270.350 to MES. For the purpose of open access, the authors have applied a Creative Commons Attribution (CC BY) license to any Author Accepted Manuscript version arising from this submission.

## Figure Legends

**Figure S1: Trimming of superfluous pixels. (A)** A branch end is recognized as a line of pixels with a single branch point and a single terminus. The binary code represents background (0) and vein (1). Note values of one are no longer present after trimming **(B).** So, if the length of this line is below the trim_factor threshold, i.e., it is small and therefore not considered a vein but an array of superfluous pixels, it is trimmed. Scale bars represent 200μm.

**Figure S2: Comparison between Starch4Kranz and LeafVeinCNN. (A)** Starch4Kranz correlated more strongly with manual traces compared with LeafVeinCNN **(B)**. Lower vein densities were detected less well by LeafVeinCNN **(B)** compared with Starch4Kranz **(D)**. r = Pearson’s correlation coefficient; LVCNN – LeafVeinCNN. Scale bars represent 200μm. Skeletons have been thickened to ensure visibility.

**Figure S3: Removal of commissural veins via pattern recognition. (A)** A commissural vein is recognized as being a pixel with three branch points. The binary code in the circle represents background (0) and vein (1). Note values of one are no longer present once commissural vein are removed **(B)**. Scale bars represent 200μm.

**Figure S4: Automated tracings for each accession of maize.** Representative outputs from Starch4Kranz on each founder from the MAGIC maize population. Scale bars represent 200μm. Skeletons have been thickened to ensure visibility.

