## Supplementary material for "A rapid method to quantify vein density in C_4_ plants using starch staining": S Figure 1-4

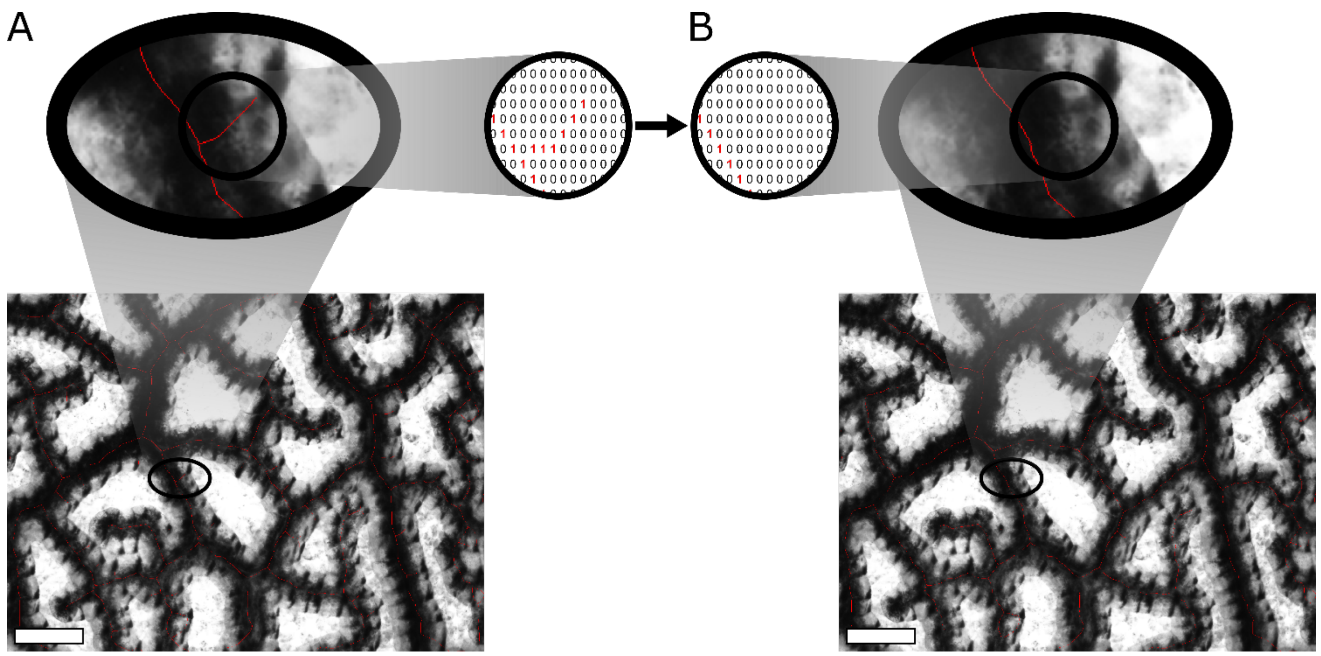


**Figure S1: Trimming of superfluous pixels. (A)** A branch end is recognized as a line of pixels with a single branch point and a single terminus. The binary code represents background (0) and vein (1). Note values of one are no longer present after trimming **(B).** So, if the length of this line is below the trim_factor threshold, i.e., it is small and therefore not considered a vein but an array of superfluous pixels, it is trimmed. Scale bars represent 200μm.

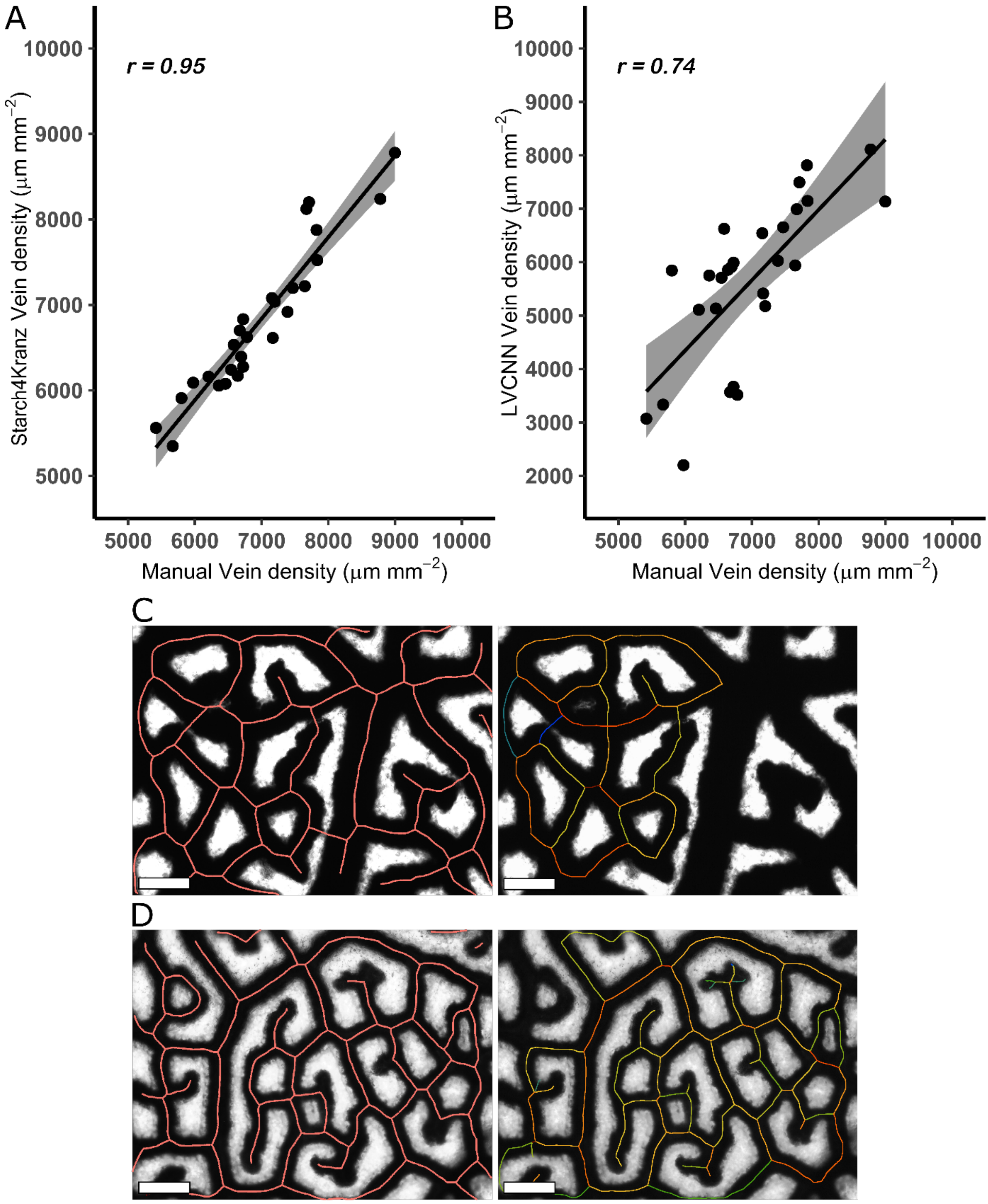


**Figure S2: Comparison between Starch4Kranz and LeafVeinCNN. (A)** Starch4Kranz correlated more strongly with manual traces compared with LeafVeinCNN **(B)**. Lower vein densities were detected less well by LeafVeinCNN **(B)** compared with Starch4Kranz **(D)**. r = Pearson's correlation coefficient; LVCNN – LeafVeinCNN. Scale bars represent 200μm. Skeletons have been thickened to ensure visibility.

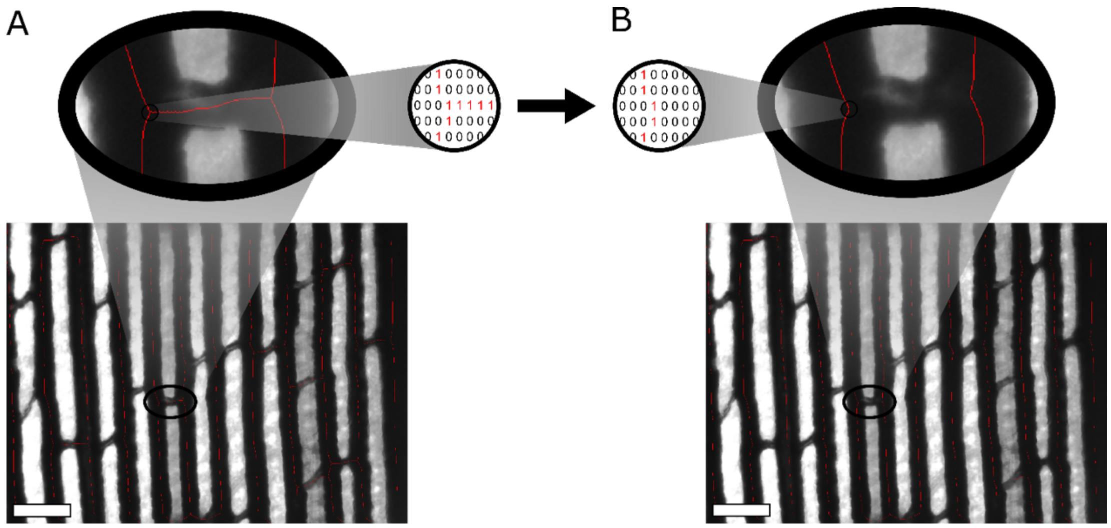


**Figure S3: Removal of commissural veins via pattern recognition. (A)** A commissural vein is recognized as being a pixel with three branch points. The binary code in the circle represents background (0) and vein (1). Note values of one are no longer present once commissural vein are removed **(B)**. Scale bars represent 200μm.

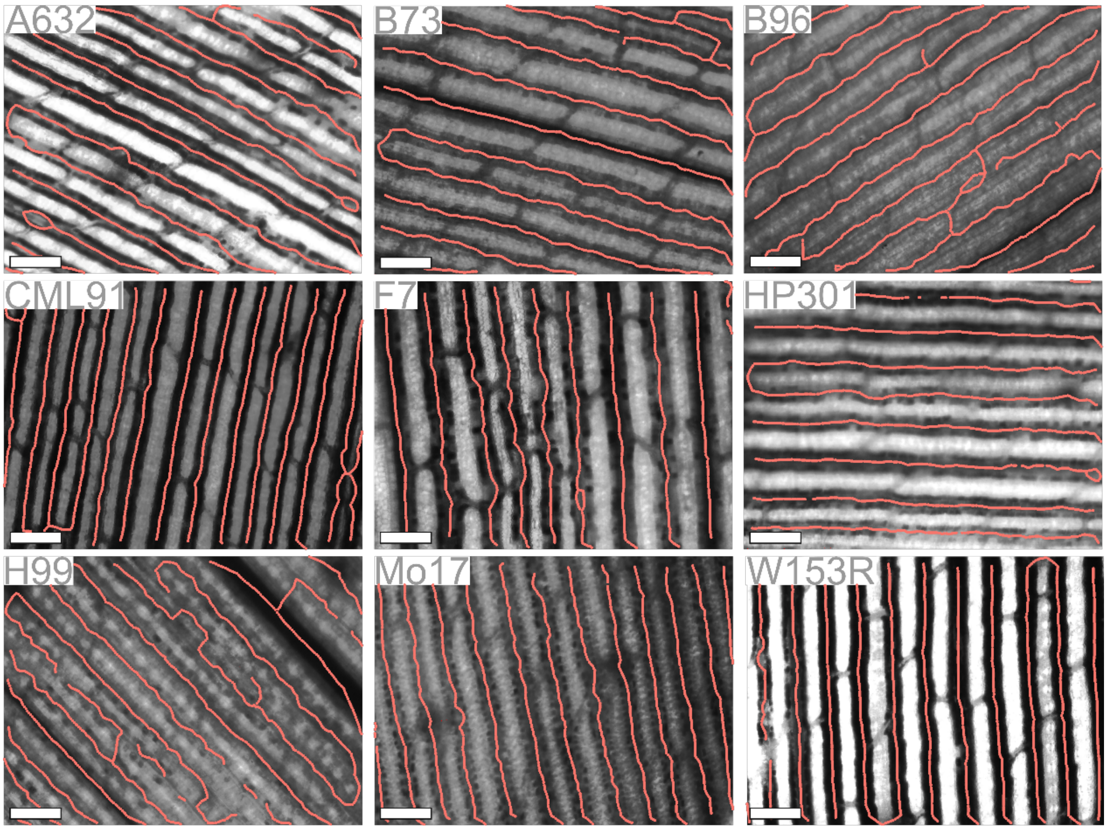


**Figure S4: Automated tracings for each accession of maize.** Representative outputs from Starch4Kranz on each founder from the MAGIC maize population. Scale bars represent 200$\mu$m. Skeletons have been thickened to ensure visibility.
